# Anti-Bacterial Activity of Substances Produced from Lactic Acid Bacteria in Metata Ayib (Traditional Ethiopian Spiced Fermented Cottage)

**DOI:** 10.1101/2020.06.19.161125

**Authors:** Alazar Essayas, Sujata Pandit, Deepak Kumar Verma

## Abstract

There have been incerases in antibiotic resistant strains of human pathogens and causing treatment of microbial infections difficult. These days more attention is given to seraching new antimicrobial drugs to combat pathogenic microbes. Traditional fermented cottage of Metata Ayib which is naturally enriched with lactic acid bacteria (LAB) and used to preserve cheese for long time in Northern Ethiopia, may have antimicrobial activity against various human pathogens. However, there was no scientific report on the antimicrobial activity of lactic acid bacteria isolated from Metata Ayib. The objective of this study is to evaluate antibacterial activity of lactic acid bacteria from *Metata Ayib* against clinical and standard human pathogens. The study was laboratory based experiment. Antibiotic production by the LAB was performed by inoculating the LAB isolates into 6.0 ml MRS medium and incubating at 30 °C. Cell free supernatants (CFS) were collected by centrifugation (10,000 rpm for 15 min at 4 °C) of the six day fermented broth cultures. The pH of the CFS was adjusted to 6.5 with 4 N NaOH to eliminate the effect of organic acids.The susceptibility of produced antibiotic against test organisms was done by growing on Muller Hinton agar in triplicate using well diffusion method and the inhibition zone was recorded. Preceddingly, MIC and MBC was determined using standard methods. The antibiotic substance exhibited antimicrobial activity towards standard and drug resistant bacteral strains with the inhibition zone ranges upto 25.33±3.21 mm. The result of MIC against tested ornaisnms showed a considerable antimicrobial activity of the antibiotic substance withn the range values 6.25% to12.5%. In addition, the result of this study showed that LAB obtained from Metata Ayib exhibits antimicrobial activity against standard and pathogenic pathogenic test bacterial species ranged from 12.5-25% of MBC value. This might be due to the production of organic acids, but also other compounds, such as ethanol, hydrogen peroxide, diacetyl, reuterin and bacteriocins. The results of this investigation can also provide baseline for information for future studies about the application of antibacterial substances produced by LAB from fermented Metata ayib.

## 1. Introduction

Microbial fermentation has played an important role in food processing for thousands of years. It provides a way to preserve food products by increasing their quality as well as safety and reducing the energy required for cooking. These significant changes develop new aroma, flavor, taste and texture that increase the palatability and the shelf life of the food. Such activities are performed by using different metabolites produced from different microorganisms like LAB (Assefa et al, 2008). Lactic acid bacteria plays an important role in the fermentation of dairy products. The fermenting species of LAB such as, *Leuconostoc, Streptococcus, Lactobacillus, Enterococcus* and *Pediococcus*. LAB confers preservative and detoxifying effects on food as well(Beresford et al., 2001). When used regularly, LAB fermented foods boost the immune system and strengthen the body in the fight against pathogenic bacterial infections. Milk is a globally known nutritious food, not only for the infants, but for human adults as well. Dairy products such as yoghurt, cheese and sour milk are fermented, protein-rich milk products. Several probiotics in use today are obtained from fermented milk products. Yoghurt, a fermented milk product, offers other benefits such as the ability to kill pathogens as well as modulate the immune system (Chelule et al., 2010). Cheese micro flora is divided into two groups. Primary group includes starter flora which refer to starter LAB and secondary group includes non starter lactic acid bacteria (NSLAB), propionic acid bacteria (PAB), smear bacteria, moulds and yeast.

Fermented milk has so many advantages than the unfermented one, it provide a way to preserve food products, to enhance the nutritional value and also to inhibit the growth of pathogenic bacteria on the milk product. The process also increases digestibility and exert health promoting benefits. Fermentation may assist in the destruction or detoxification of certain undesirable compounds which may be present in raw foods. These are compounds such as phytates, polyphenols and tannins (Das and Deka, 2012).

The microorganisms behind this milk fermentation are LAB. LAB are group of gram positive, non spore forming, non–respiring, cocci or rods, which produce lactic acid as the major end product of the fermentation of carbohydrate. They produce lactic acid as a result of carbohydrate fermentation.

LAB is Widely used as starter cultures in the manufacture of fermented products including milk products such as yoghurt, cheese, meat products, bakery products wine and vegetable. They are used as natural or selected starters in food fermentation in which they perform acidification due to production of lactic acid flavor. Bacteria from genera *Lactobacillus, Lactococcus, Leuconostoc, Pediococcus* and *Streptococcus* are the main species of LAB involved. Although several more have been identified but they play a minor role in lactic fermentations. LAB growth lowers both the carbohydrate content of the food that they ferment and the pH due to the lactic acid production. LAB are widespread in nature, their nutritional requirements are very complex. Hence, they predominate habitats that rich in carbohydrates, protein breakdown products, vitamins and environments with low oxygen. This confirms the prevalence in dairy products (Stiles and Holzapfel, 1997).They are generally associated with habitat rich in nutrients such as milk, cheese, meat, beverages and vegetables. They could be also isolated from soil, lakes, intestinal tract of animals and humans. LAB are often inhibitory to other microorganisms and this is the basis of their ability to affect the keeping quality and safety of many food products. Many of LAB species received a generally recognized as safe (GRAS) or qualified presumption of safety (QPS) -status. LAB is traditionally been associated with these dairy products and with cereal-, vegetable- and meat-based fermented foods, either as intentionally added starters or due to their natural presence leading to spontaneous fermentation. Certain LAB are also used as probiotics added to confer health benefits to consumers or to anti microbial activity.

Metata Ayib can stay for a long storage time unlike the Ayib which has few days shelf life (Assefa, et al, 2008). Even though using Metata Ayib is widely practiced in the north west Ethiopia, scientific research that address the anti microbial activity of LAB from Metata Ayib has not been well conducted yet. Therefore the knowledge of the antimicrobial activity of LAB from Metata Ayib can help to provide basic information necessary to understand the role of LAB as an anti microbial effect. The outcome of this study may help to have a brief understanding about the anti bacterial activity of antibiotic substances from Metata ayib against common pathogenic bacteria.

## 2. LITERATURE REVIEW

### 2.1. Traditional Fermented Dairy Products in Ethiopia

Ethiopians have a culture of using milk in different types fermentative forms. The fermented product has different domestic names such as *ergo*, *ititu*, *geinto* or Metata among the Amhara, Oromo, Sidama, or Wolayta people, respectively(Ashenafi and Mehari, 1995). The fermentation is usually natural. LAB also become established on the inner walls of the container and serve as starter culture. In most cases, this is made possible through the proliferation of the initial milk flora, with microbial succession determined by ambient temperatures and chemical changes in the fermenting milk. (Anteneh, et al., 2011).

Raw milk is left either at ambient temperatures or kept in a warmer place to ferment. In rural areas, particularly among the pastoralists, raw milk is usually kept in a well-smoked container and milk from a previous fermentation serves as source of inoculums. Incubation temperature does not usually vary significantly, particularly in the lowlands, and the taste of the fermented product may, in general, be more or less uniform. The fermented product may also be processed into traditional butter (*qibe*) and butter milk (*arrera*)(Ashenafi and Beyene, 1993)

### 2.2. Ayib (Ethiopian Cottage Cheese)

In Ethiopia, milk processing is based on sour milk mainly due to high ambient temperatures, consumer’s preference and increased keeping quality of sour milk (O’Mahony, et al., 1988). *Ayib* is a traditional Ethiopian cottage cheese made from sour milk after the fat is removed by churning and produced by slowly heating naturally soured milk until a distinct curd mass forms and floats over the whey. The whey is traditionally known as *aguat*. It is an acidic product. *Ayib* is a popular milk product consumed by the various ethnic groups in the country(Anteneh, et al., 2011). It is an important source of nutrients and serves as a staple diet. It may be consumed fresh as side dish, or it may be spiced with hot spices, salt and other herbs (Gonfa, et al., 2001). Raw milk is collected in a clay pot and kept in a warm place (about 30°C) for 24 to 48 hours to sour spontaneously. The pH of sour milk is usually about 4. Churning of sour milk is carried out by slowly shaking the contents of the pot until the fat is separated. Temperature of *ayib* preparation could vary between 40°C to 70°C without any significant impact on composition and yield.

However, temperature above 80°C imparts a cooked flavor to the product. Cooking at 60°C markedly decreased the number of most bacterial groups, yeasts and molds. Heat treatment at 70°C required a relatively shorter time for curd formation and achieved maximum reduction in number of the various microbial groups. Since temperatures higher than 80°C are reported to give the product a cooked flavor(Anteneh et al., 2011). Heat precipitation of curd at 70°C (pH 4.0) was recommended as it resulted in a less contaminated and more wholesome *ayib*(Anteneh, et al., 2011). After gradual cooling, the curd is recovered from the whey. *Ayib* comprises 79% water, 14.7% protein, 1.8% fat, 0.9% ash and 3.1% soluble milk constituents and the yield should be at least 1 kg of *ayib* from 8 liters of milk (12.5%)(O’mahony, et al., 1988). Fekadu and Abrahamsen (2016) analyzed various ayib samples produced by small-holders in three regions of southern Ethiopia and found out that the samples consisted of 80 - 81% moisture, 13.4 - 16% protein, 1.9 - 2.0% fat and 0.75 - 0.87% minerals (Gebreselassie et al., 2016).

### 2.3. Factors Influencing the Growth of Microorganisms in Cheese

The survival and growth of pathogens in cheese depend on the many factors, including variations in pH, salts, the presence of competing microbiota, and the temperature and biochemical changes during ripening. The microbiological quality of the milk and the good manufacturing practices will also contribute to the safety of the final product; especially in cheeses where milk is not pasteurized (Beresford et al., 2001).

### 2.4. Metata Ayib

*Metata ayib* is a traditional fermented cottage cheese produced in Northwest Ethiopia. It differs from the traditional cottage cheese *Aiyb* in that its production involves the use of different spices and spontaneous fermentation for 20 days. *Metata ayib* has a long shelf life up to one year in semi-solid form but more than ten years in dry form. In contrast *Ayib* has a shelf life of only a few days. The property of *Metata ayib* has not been fully understood and characterized(Seifu, 2013).

### 2.5. Lactic Acid Bacteria

LAB are gram-positive, acid -tolerant and non-spore forming cocci and rods. They are a heterogeneous group of bacteria comprising about 20 genera within the phylum Firmicutes. From a practical point of view,the genera *Aerococcus*, *Carnobacterium*, *Enterococcus*, *Lactobacillus*, *Lactococcus*, *Leuconostoc*, *Oenococcus*, *Pediococcus*, *Streptococcus*, *Tetragenococcus*, *Vagococcus* and *Weissella* have been considered as the principal LAB (Holzapfel et al., 2001).

One common feature of the LAB is their ability to produce lactic acid as a major end product of their fermentation of hexoses. As fermenting organisms, LAB lack electron transport systems and cytochromes, and they do not have a functional Krebs cycle. Based on end products of glucose metabolism, LAB can be divided into two groups, namely homofermentative and heterofermentative(Jay, 1998). Hetrofermenttive Organisms metabolize glucose via the pentose phosphate pathway. End products can vary depending upon level of aeration and presence of other proton and electron acceptors. Acetyl-phosphate can be converted to acetate and ATP or reduced to ethanol without ATP production (Anteneh, et al. 2011).

LAB are widespread organisms and they may be found in many environments rich in carbohydrates. In addition to carbohydrates, LAB have complex nutritional requirements for aminoacids, peptides, fatty acid esters, salts, nucleic acid derivatives and vitamins In short, they have complex nutritional requirements due to their lack of many biosynthetic pathways. On the other hand, they are found in a wide range of different environmental niches due to their good capacity for adaptation. In food products, they are found in dairy products, such as yoghurt and cheese, in fermented vegetables, in fermented meats and in sourdough bread(Tannock, 2004).

**Figure 1.**
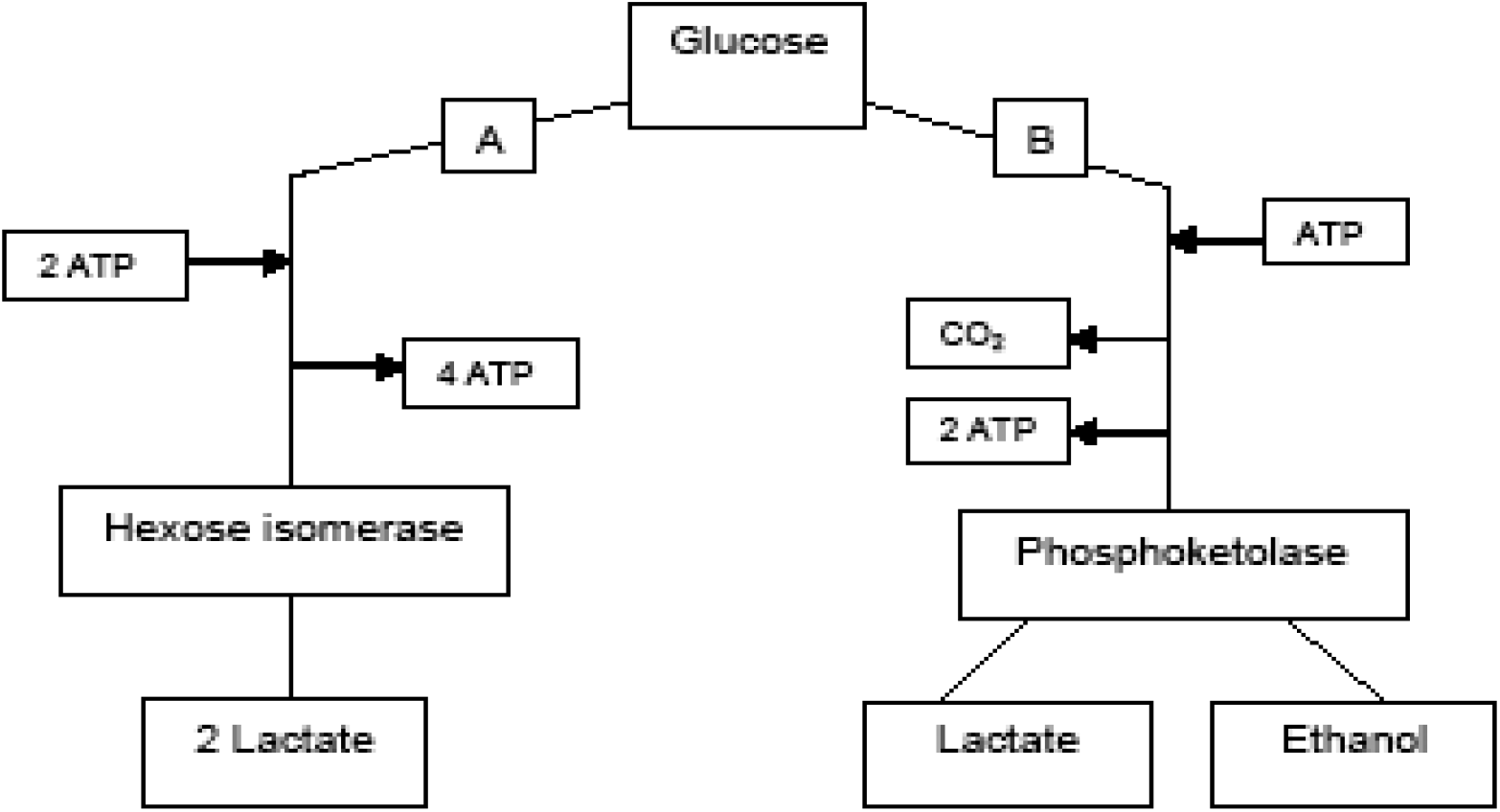
Generalized pathways for the production of fermentation products from glucose by A: homofermentative LAB and B: heterofermentative LAB.

### 2.6. Classification of LAB

The general basis for the classification of bacteria is based on international agreement.This classification system at genus level first divides the LAB according to morphology. The next important characteristics used in differentiation of LAB genera is the mode of glucose fermentation under standard condition (non limited supply with glucose,growth factors such as amino acids,vitamins and nucleic acids precursors and limited oxygen availability, under this condition, LAB can be divided in to two groups, homofermentative and hetrofermentative. homofermentative LAB convert sugars almost quantitie to lactic acid, the second group hetrofermentative bacteria produce not only lactic acid but ethanol acetic acids and carbon dioxide.

### 2.7. Antibacterial activity of LAB

LAB display a wide range of antimicrobial activities. Amongst these activities, the production of lactic acid and acetic acid is obviously the most important. However, certain strains of LAB are further known to produce bioactive molecules such as ethanol, formic acid, fatty acids, hydrogen peroxide, diacetyl, reuterin, and reutericyclin. Many strains also produce bacteriocins and bacteriocins-like molecules that display antibacterial activity. Besides the production of bacteriocins, some LAB are able to synthesize other antimicrobial peptides that may also contribute to food preservation and safety. For instance, strains of *Lactobacillus plantarum*, isolated from sourdough and grass silage, display antifungal activity, due to the production of organic acids, other low-molecular-mass metabolites, and/or cyclic dipeptides(Israti, D., 2011). Studies of antibiotic resistance of LAB have not been extensively investigated until recently, in contrast to the situation with pathogenic species and their antibiotic resistance. However, interest in LAB and their antibiotic resistances has recently gained attention since the resistant determinants are known to be able to be transferred between bacterial species, also from beneficial bacteria to pathogens. The first step in characterizing LAB species as being either susceptible or resistant to antibiotics is to determine the susceptible/resistant patterns with phenotypic methods. LAB give fermented milk the slightly sharp and sour taste. Additional characteristic flavor and aroma are often the result of other products of LAB. For example, acetaldehyde is known to provide the characteristic aroma of yoghurt while diacetyl imparts a buttery taste to other fermented milks. Inhibition activity of LAB has been reported to be due to a combination of many factors such as production of lactic acid which brings about reduction of pH of the fermentation medium and production of inhibitory bioactive compounds such as hydrogen peroxide and bacteriocins which are responsible for most antimicrobial activity. LAB play a major part in most fermentation processes, not only because of their ability to improve the flavor and aroma but especially for their preservative effects on food(Egerva et al., 2007).

Species belong to *Streptococcus* and *Leuconostoc* produce the least amount of acid; while the homofermentative species of *Lactobacillus* produce the greatest amount of acid. Hetrofermentative *Leuconostoc* and *Lactobacillus* species convert glucose to about 50% lactic acid, 25% acetic acid and ethyl alcohol, and 25% carbon dioxide. This is important in flavour development and in leavening of certain bread fermented foods like bread(Jay, 1998).

### 2.8. Antimicrobial Compounds Produced by Lactic Acid Bacteria

The preservative action of starter culture in food and beverage systems is attributed to the combined action of a range of antimicrobial metabolites produced during the fermentation process. These include many organic acids such as lactic, acetic and propionic acids produced as end products which provide an acidic environment which creates unfavourable condition for the growth of many pathogenic and spoilage microorganisms (Yvon and Rijnen, 2001). Acids are generally thought to exert their antimicrobial effect by interfering with the maintenance of cell membrane potential, inhibiting active transport, reducing intracellular pH and inhibiting a variety of metabolic functions (Daeschel, 1993).

Acidic metabolic produts have a very broad mode of action and inhibit both gram-positive and gram-negative bacteria as well as yeast and moulds. One good example is propionic acid produced by propionic acid bacteria, which has formed the basis for some biopreservative products, given its antimicrobial action against microorganisms including yeast and moulds(Israti, D., 2011). In addition to acids, starter strains can produce a range of other antimicrobial metabolites such as ethanol from the hetrofermentative pathway, H_2_O_2_ produced during aerobic growth and diacetyl which is generated from excess pyruvate coming from citrate. In particular H_2_O_2_ can have a strong oxidizing effect on membrane lipids and cellular proteins and is produced using such enzymes as the flavor protein oxidoreductases NADH peroxidase, NADH oxidase and glycerophosphate oxidase(Daeschel, 1993).Obviously, each antimicrobial compound produced during fermentation of milk produts provides an additional hurdle for pathogens and spoilage bacteria to overcome before they can survive and/or proliferate in a food or beverage, from time of manufacture to time of consumption. Since any microorganism may produce a number of inhibitory substances, its antimicrobial potential is defined by the collective action of its metabolic products on undesirable bacteria(Israti, D., 2011). Other examples of secondary metabolites produced by LAB which have antagonistic activity include the compound reuterin and the recently discovered antibiotic reuterocyclin, both of which are produced by strains of *Lactobacillus reuteri*. Reuterin is an equilibrium mixture of monomeric, hydrated monomeric and cyclic dimeric forms of hydroxypropionaldehyde (Anteneh,et al., 2011).

It has broad spectrum of activity and inhibits fungi, protozoa and a wide range of bacteria including both gram-positive and gram-negative bacteria. This compound is produced by stationary phase cultures during anaerobic growth on a mixture of glucose and glycerol or glyceraldehydes. Consequently, in order to use reuterin-producing *L. reuteri* for biopreservation in a food product, it would be beneficial to include glycerol with the strain(Israti, D., 2011). Results demonstrated that after 6-day of storage, there were approximately 100-fold-less gram negative bacteria in the *L. reuteri* samples than in the untreated control. More recently, the first antibiotic produced by a LAB was discovered. Reuterocyclin is a negatively charged, highly hydrophobic antagonist, and structural elucidation revealed it to be a novel tetramic acidd(Daeschel, 1993).

The spectrum of inhibition of the antibiotic is confined to gram-positive bacteria including *Lactobacillus* spp.*, Bacillus subtilis, Bacillus cereus, Enterococcus faecalis, Staphylococcus aureus* and *Listeria innocua*. Interestingly, inhibition of *Escherichia coli* and *Salmonella* is observed under conditions that disrupt the outer membrane, including truncated lipopolysaccharides (LPS), low pH and high salt concentrations(Israti, D., 2011). Since it is well known that nisin can kill gram-negative bacteria under conditions which disturb the outer membrane, it is likely that there are similarities in the mode of action of nisin and this novel antibiotic. (Yvon and Rijnen, 2001).

## 3. MATERIAL AND METHOD

### 3.1. Description of the Study Area

The study was conducted at the University of Gondar, North Gondar of Amhara Region which is located at about 723 kms away from Addis Ababa, the capital city of Ethiopia. The university is geographically located at an altitude of around 2,200 m (above sea level), 12° 36’ 0” of latitude and 37° 28’ 0” of longitude and has a reliable rainfall averaging between 800 and 1000 mm per annum

### 3.2. Study Design

Laboratory based experiment was conducted to identify the antimicrobial activity of LAB from *Metata ayib* (Ethiopian spiced traditionally fermented cheese).

### 3.3. Production of Antibiotic Substances

Microorganisms used for the production of antibiotic substances were obtained from Biotechnology laboratory, previously identified and isolated from Metata Ayib. The organisms were characterized up to the genius level by Tsehayneh Geremew, during his MSC thesis work. They were represented as A, B, C, D, E and F, for *Lactobacillus, Lactococcus, Streptococcus, Pediococcus, Weissella* and *Entrococcus, respectively*. Antibiotic production by the LAB was performed by inoculating the LAB isolates into 6.0 ml MRS broth medium and incubating at 30 °C. Cell free supernatants (CFS) were collected by centrifugation (10,000 rpm for 15 min at 4 °C) of the overnight broth cultures. The pH of the CFS was adjusted to 6.5 with 4 N NaOH to eliminate the effect of organic acids. Possible inhibition by the hydrogen peroxide was also removed by the addition of catalase at a final concentration of 1.0 mg/ml at 30 °C for 1.

### 3.4. Sources of Test Organisms and Inoculum Preparation

In this study,standard and drug resistance pathogenic bacteria were used for determination of antibacterial activites of the antiibiotic susbstance from Metata ayib.In brief the bacterial strains used to asses the anti bacterial properties of the antibiotic substance include both gram posetive and gram negative strains: *S. aureus* (ATCC25923), *S. pneumoniae* (ATCC49619), *S. flexneri* (ATCC 12022), *E. coli* (ATCC25922) were the standard bacterias used,Multidrug resistant test bacterias were:K.pneumoniae (drug resistant), *S. pneumoniae(drug resistant), E.coli(drug resistant)*. The microorganisms were selected because they were of great public health importance. The organisms were obtained from the Department of Medical Microbiology, University of Gondar.

The antibacterial activity was tested at Molecular Biology Laboratory Department of Biotechnology, University of Gondar. Standard pathogenic bacteria were inoculated and spread over on Muller-Hinton (MH) agar (HIMEDIA, India) and incubated for 24 h. From 2-3 colonies were picked up by wire loop aseptically into sterile normal saline solution and the turbidity were adjusted to 0.5 McFarland’s standard solution (a concentration of 1.5 × 10^8^ CFU/ml).

### 3.5. Preparation of 0.5 McFarland turbidity standards

The McFarland 0.5 turbidity standard was prepared by adding 50 μl of a 1.175% (wt/v) barium chloride dehydrate (BaCl_2_.2H_2_O) solution to 9.95 ml of 1% (v/v) sulfuric acid. McFarland standard tube were then sealed with paraffin to prevent evaporation and stored in dark at room temperature. The accuracy of the density of a prepared McFarland standard was checked using a spectrophotometer with a 1-cm light path length. For the 0.5 McFarland standards, the absorbance was adjusted at a wavelength of 625nm and water was used as a blank solution. The 0.5 McFarland standards were vigorously agitated to turbidity on a vortex mixer before use. As with the barium sulfate standards, a 0.5 McFarland standard is comparable to a bacterial suspension of 1.5×10^8^ colony-forming units (CFU)/ml (NCCLS, 2000).

### 3.6. Susceptibility Test

The agar-well diffusion method prescribed by NCCLS (2000) was employed in the susceptibility testing. Suspensions of the standard and drug resistnat test bacteria were made in sterile normal saline solution and adjusted to the 0.5 McFarland’s standard. Each Mueller Hinton (MH) agar plate was uniformly seeded with 0.1 ml of the respective test organism and spread with sterilized cotton swab. The plates were left on the bench for excess fluid to be absorbed. Wells of 6 mm in diameter, 5 mm deep and about 2 cm apart were punched in the MH agar with a sterile cork borer. Then 100 μl of the antibiotics extracts were dropped into each well till its fullnes. Inoculated plates were incubated at 37°C over night. After a 24-hour incubation period, the mean zones of inhibition were thereafter measured in mm, for the entire individual test organism. Distilled water was used as a negative control, whereas vancomycine (30μg) was used as a positive control. The experiment was performed in triplicates and the interpretation of antibacterial properties was conducted according to CLSI statndard. Inhibition zones greater than 15 mm were categorized as strong activity, from 10-15 mm as moderate activity and less than 10 mm as weak activity.

### 3.7. Determination of the Minimum Inhibitory Concentrations(MIC)

The minimal inhibitory concentration of were determined based using macro-test tube dilution method. In the determination of MIC,30 sterile screw capped test tubes were placed on suitable rack in three rows i.e., 10 test tubes in each row and labeled each of them including the negative and positive control test tubes. The antibiotic substance was diluted to concentrations (50%, 25%, 12.5%, 6.25%). The first test tube contained 50% of the extract (4 ml of extract and 4 ml of nutrient broth) the second one contained 25% extract (2ml of extract and 6 ml of nutrient broth) the third one contained 12.5% extract (1 ml of extract and 7 ml of nutrient broth)and the last one contained 6.25% of extract(0.5 ml of extract and 7.5 ml of nutrient broth) to bring 8 ml.

To each of dilution, 100 micro liter of bacterial inoculums was carefully dispensed in sterile screw capped tubes which consists antibiotic substance and nutrient broth. Nutrient broth with bacterial inoculation but no any extract(positive control tubes) and nutrient broth only with no bacterial inoculation (negative control tubes) were included for every test organism to demonstrate an adequate microbial growth over the course of the incubation period and media sterility, respectively. Then tubes were incubated aerobically at 37 °C for 24 hours and examined for bacterial growth. The lowest concentration (highest dilution) of the extract and that produced no visible bacterial growth (turbidity) after overnight incubation was recorded as MIC.

### 3.8. Determination Of Minimum Bactericidal Concentration(MBC)

To determine the MBC,macro test tubes used in the MIC, which did not show any visible growth of bacteria after the incubation period were sub cultured on to the surface of the freshly prepared Mueller Hinton Agar (MHA) plates and incubated at 37 °C for 24 hours. The MBC was recorded as the lowest concentration (highest dilution) of the extract that did not permit any visible bacterial colony growth on the agar plate after the period of incubation (Mueller and Mechler, 2008).

### 3.9. Statistical Analysis

All statistical analyses were conducted using SPSS software version 20.0. All the experiments were conducted in triplicate and the different data were analyzed and compared statistically using ANOVA at 95% level of significance. A probability value at p ≤ 0.05 was considered statistcically significant. Data are presented as mean values ± standard deviation calculated from triplicate determinations.

## 4. RESULT

In this investigation, antibiotic substances were produced from iolate A, B, C, D, E and F represented for *Lactobacillus, Lactococcus, Pediococcus, Streptococci, Weissella* and *Entrocossus* respectively. Their antibiotic activitites were determined using standard methods aganist some standard and multidrug resistant pathogenic bacteria. The antibiotic activity of antimimicrobial substances produced from isolate A aginst standard and multidrug resistant pathogenic bacteria was presented on Table 1A. The inhibition zone of antibiotic substances (18.00±2.00) from isolate A against *S. flexneri* (ATCC 12022) was not stastistically (P ≥ 0.05) different from inhibition zone of vacomycin (VM) (20.00±2.00). Similarly, the inhibition zone of antibiotic substances (13.33±1.15) from isolate A a against *E. coli* (ATCC25922) was almost stastistically (P ≥ 0.05) similar with the inhibition zone of VM. These two pathogenic bacteria are standard pathogens (sensitive for most antibiotics). The inhibition zone of antibiotic substances produced by isolate A (12.66±1.15) aginst *K. pneumonae* (multidrug resistant strain) was not significantly (P ≥ 0.05) different from the inhibition zone of VM (16.00 ± 1.73). Surprisingly the inhibition zone of antibiotic substances (18.00±2.00) produced by isolate A to *MRSA* (multidrug resistant strain) was significantly (P ≥ 0.05) greater than the inhibition zone of VM.

**Table 1A:**
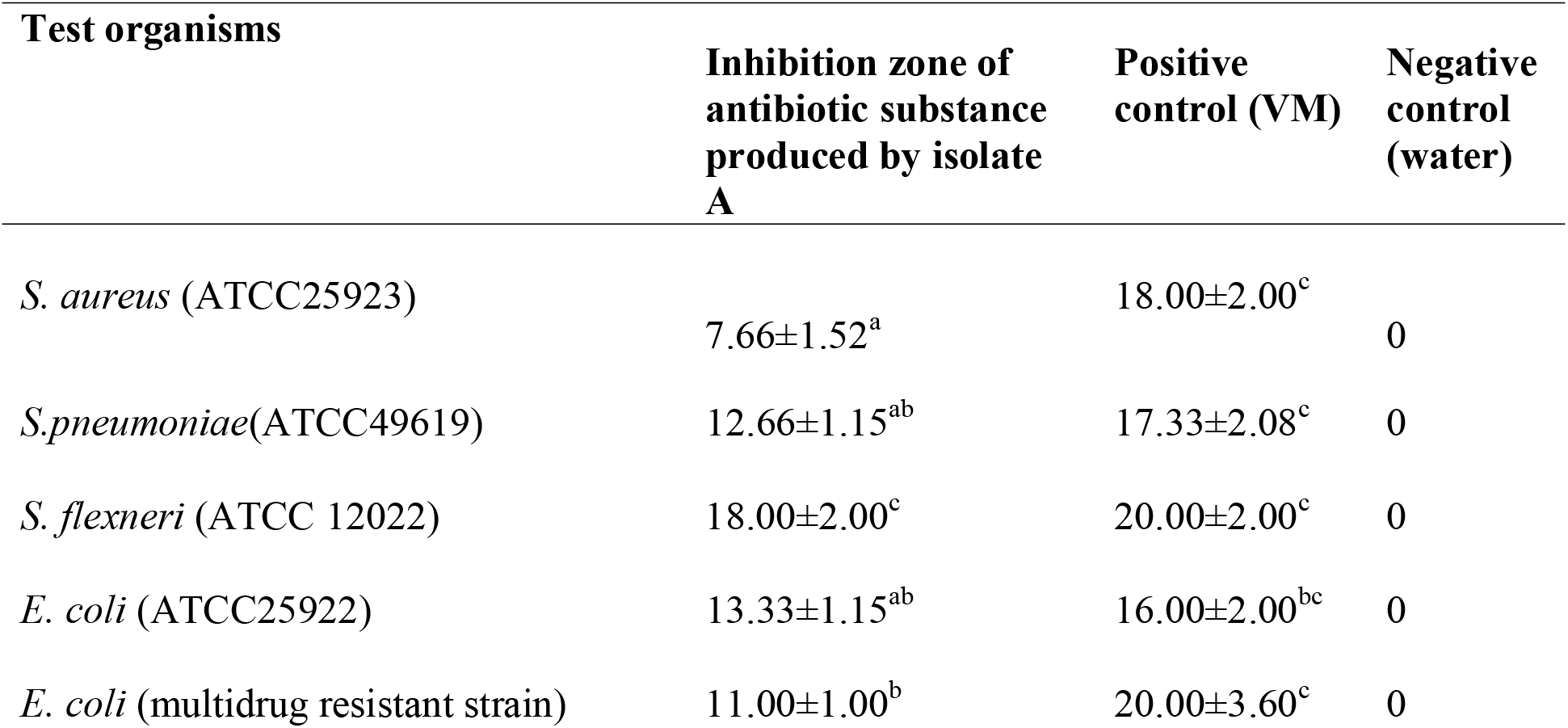
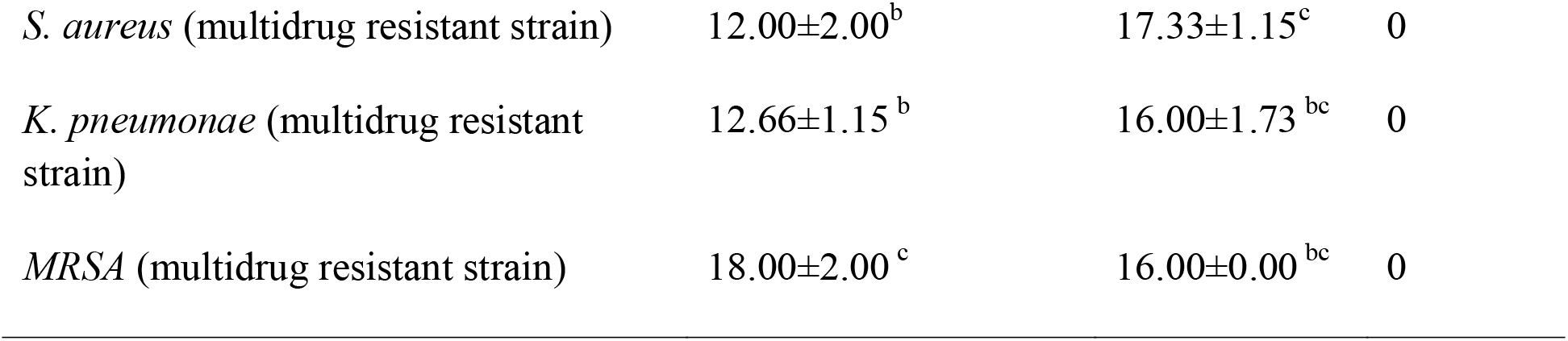
The mean inhibition zone of antibiotic substance produced by LAB against standard and drug resistant clinical pathoegnic bacteria

| Test organisms | Inhibition zone of antibiotic substance produced by isolate A | Positive control (VM) | Negative control (water) |
| --- | --- | --- | --- |
| *S. aureus* (ATCC25923) | 7.66±1.52<sup>a</sup> | 18.00±2.00<sup>c</sup> | 0 |
| *S.pneumoniae*(ATCC49619) | 12.66±1.15<sup>ab</sup> | 17.33±2.08<sup>c</sup> | 0 |
| *S. flexneri* (ATCC 12022) | 18.00±2.00<sup>c</sup> | 20.00±2.00<sup>c</sup> | 0 |
| *E. coli* (ATCC25922) | 13.33±1.15<sup>ab</sup> | 16.00±2.00<sup>bc</sup> | 0 |
| *E. coli* (multidrug resistant strain) | 11.00±1.00<sup>b</sup> | 20.00±3.60<sup>c</sup> | 0 |
| *S. aureus* (multidrug resistant strain) | 12.00±2.00<sup>b</sup> | 17.33±1.15<sup>c</sup> | 0 |
| *K. pneumoniae* (multidrug resistant strain) | 12.66±1.15<sup>b</sup> | 16.00±1.73<sup>bc</sup> | 0 |
| *MRSA* (multidrug resistant strain) | 18.00±2.00<sup>c</sup> | 16.00±0.00<sup>bc</sup> | 0 |

Values are means of triplicate determinations; Values within the same row followed by different superscripts are significantly different at (p < 0.05).

The antibiotic activity of antimimicrobial substances produced from isolate A aginst standard and multidrug resistant pathogenic bacteria was shown on Table 1B. The inhibition zone of antibiotic substance produced by isolate B against all stansard pathogenic bacteria were siginificantly (P ≤ 0.05) less than the inhibition zone of VM. The inhibition zone of antibiotic substance produced by isolate B against *E. coli* (multidrug resistant strain), *S. aureus* (multidrug resistant strain) and *MRSA* (multidrug resistant strain) were not significantly (P ≥ 0.05) different from the inhibion zone of VM. But The inhibition zone of antibiotic substance (21.33±3.05) produced by isolate B against *K. pneumonae* (multidrug resistant strain) was stastistically (P ≤ 0.05) greater than the inhibition zone of of VM.

**Table 1B:**
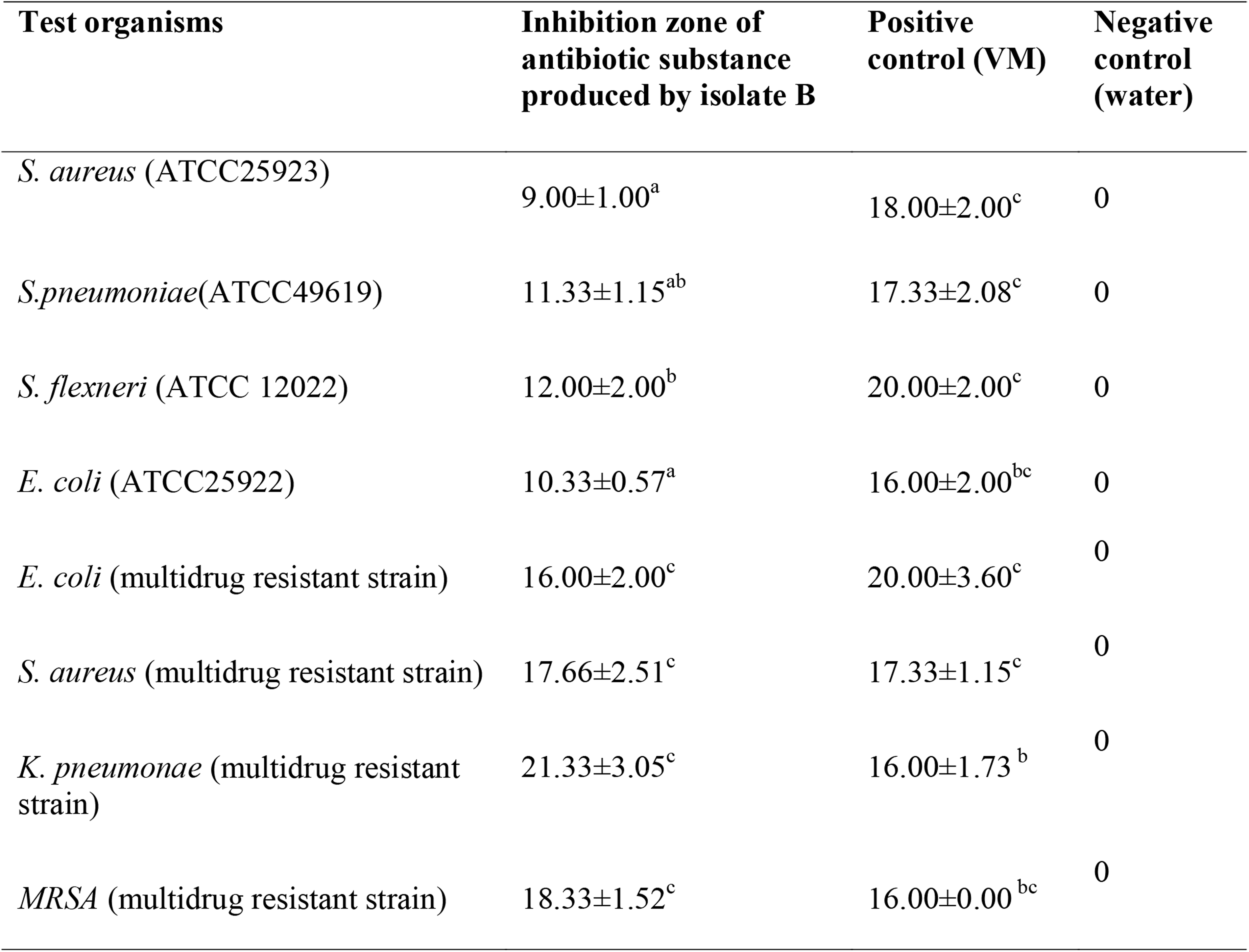
The mean inhibition zone of antibiotic substance produced by LAB against standard and drug resistant clinical pathoegnic bacteria.

| Test organisms | Inhibition zone of antibiotic substance produced by isolate B | Positive control (VM) | Negative control (water) |
| --- | --- | --- | --- |
| *S. aureus* (ATCC25923) | 9.00±1.00<sup>a</sup> | 18.00±2.00<sup>c</sup> | 0 |
| *S.pneumoniae*(ATCC49619) | 11.33±1.15<sup>ab</sup> | 17.33±2.08<sup>c</sup> | 0 |
| *S. flexneri* (ATCC 12022) | 12.00±2.00<sup>b</sup> | 20.00±2.00<sup>c</sup> | 0 |
| *E. coli* (ATCC25922) | 10.33±0.57<sup>a</sup> | 16.00±2.00<sup>bc</sup> | 0 |
| *E. coli* (multidrug resistant strain) | 16.00±2.00<sup>c</sup> | 20.00±3.60<sup>c</sup> | 0 |
| *S. aureus* (multidrug resistant strain) | 17.66±2.51<sup>c</sup> | 17.33±1.15<sup>c</sup> | 0 |
| *K. pneumoniae* (multidrug resistant strain) | 21.33±3.05<sup>c</sup> | 16.00±1.73<sup>b</sup> | 0 |
| *MRSA* (multidrug resistant strain) | 18.33±1.52<sup>c</sup> | 16.00±0.00<sup>bc</sup> | 0 |

Values are means of triplicate determinations; Values within the same row followed by different superscripts are significantly different at (p < 0.05).

Antimicrobial activity of antibiotic substance produced by isolate C is shown on Table 1C. The inhibition zone of antibiotic substances obtained from isolate C against all standard pathogenic bacteria were significantly (P ≤ 0.05) less than the inhibition zone of VM. The inhibition zone of antibiotic substances obtained from isolate C against *S. aureus* (multidrug resistant strain) and *K. pneumonae* (multidrug resistant strain) were stastistically (P ≥ 0.05) similar with the inhibione zone of VM. In this study, the interesting finding is that inhibition zone of antibiotic substances (25.33±4.16) produced from isolate C against *MRSA* (multidrug resistant strain) was significantly (P ≤ 0.05) greater than the inhibition zone of VM (16.00±0.00).

**Table 1C:** The mean inhibition zone of antibiotic substance produced by LAB against standard and drug resistant clinical pathoegnic bacteria.

| Test organisms | Inhibition zone of antibiotic substance produced by isolate C | Positive control (VM) | Negative control (water) |
| --- | --- | --- | --- |
| *S. aureus* (ATCC25923) | 10.66±1.52<sup>a</sup> | 18.00±2.00<sup>c</sup> | 0 |
| *S.pneumoniae*(ATCC49619) | 10.00±0.00<sup>a</sup> | 17.33±2.08<sup>c</sup> | 0 |
| *S. flexneri* (ATCC 12022) | 14.00±2.00<sup>ab</sup> | 20.00±2.00<sup>c</sup> | 0 |
| *E. coli* (ATCC25922) | 12.33±2.51<sup>a</sup> | 16.00±2.00<sup>bc</sup> | 0 |
| *E. coli* (multidrug resistant strain) | 12.33±0.57<sup>a</sup> | 20.00±3.60<sup>c</sup> | 0 |
| *S. aureus* (multidrug resistant strain) | 17.33±1.15<sup>c</sup> | 17.33±1.15<sup>c</sup> | 0 |
| *K. pneumoniae* (multidrug resistant strain) | 17.66±2.51<sup>c</sup> | 16.00±1.73<sup>bc</sup> | 0 |
| *MRSA* (multidrug resistant strain) | 25.33±4.16<sup>c</sup> | 16.00±0.00<sup>b</sup> | 0 |

Values are means of triplicate determinations; Values within the same row followed by different superscripts are significantly different at (p < 0.05).

Antimicrobial activity of antibiotic substance of isolate D is presented on Table 1 D. The inhibition zone of antibiotic substances obtained from isolate D against all standard pathogenic bacteria were significantly (P ≤ 0.05) less than the inhibition zone of VM. On the other hand, the inhibition zones of antibiotic substances obtained from isolate D against *S. aureus* (multidrug resistant strain) and *K. pneumonae* (multidrug resistant strain) were not stastistically (P ≥ 0.05) different from the inhibition zone of VM. However, the inhibition zones of antibiotic substances (25.33±4.16) obtained from isolate D against *MRSA* (multidrug resistant strain) was significantly (P ≤ 0.05) greater than the inhibition zones of VM (16.00±0.00).

**Table 1D:**
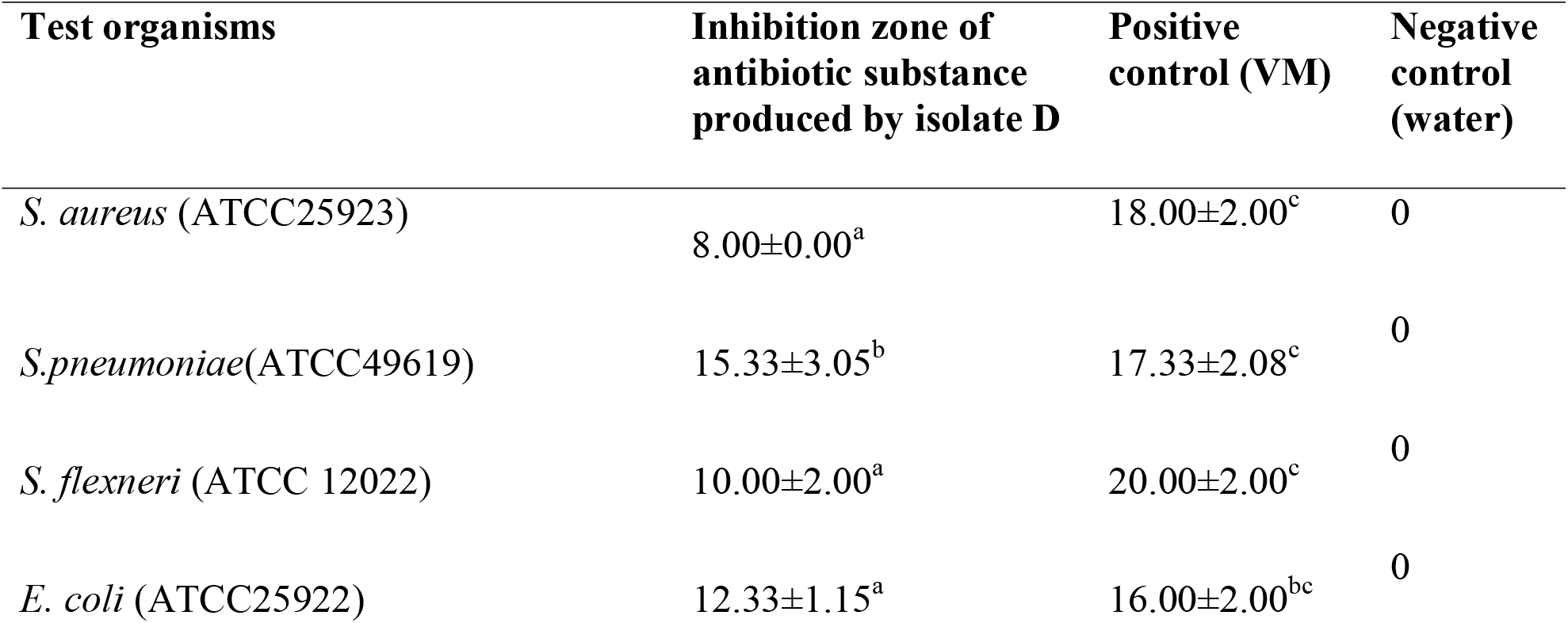
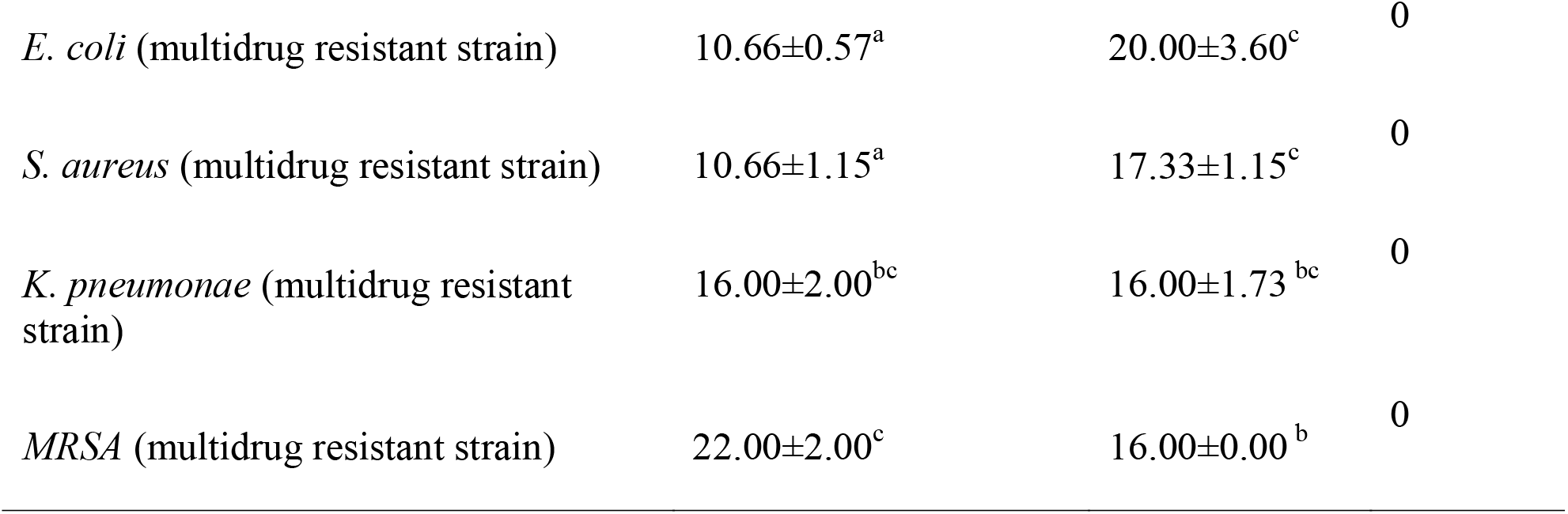
The mean inhibition zone of antibiotic substance produced by LAB against standard and drug resistant clinical pathoegnic bacteria.

| Test organisms | Inhibition zone of antibiotic substance produced by isolate D | Positive control (VM) | Negative control (water) |
| --- | --- | --- | --- |
| *S. aureus* (ATCC25923) | 8.00±0.00<sup>a</sup> | 18.00±2.00<sup>c</sup> | 0 |
| *S.pneumoniae*(ATCC49619) | 15.33±3.05<sup>b</sup> | 17.33±2.08<sup>c</sup> | 0 |
| *S. flexneri* (ATCC 12022) | 10.00±2.00<sup>a</sup> | 20.00±2.00<sup>c</sup> | 0 |
| *E. coli* (ATCC25922) | 12.33±1.15<sup>a</sup> | 16.00±2.00<sup>bc</sup> | 0 |
| *E. coli* (multidrug resistant strain) | 10.66±0.57<sup>a</sup> | 20.00±3.60<sup>c</sup> | 0 |
| *S. aureus* (multidrug resistant strain) | 10.66±1.15<sup>a</sup> | 17.33±1.15<sup>c</sup> | 0 |
| *K. pneumoniae* (multidrug resistant strain) | 16.00±2.00<sup>bc</sup> | 16.00±1.73<sup>bc</sup> | 0 |
| *MRSA* (multidrug resistant strain) | 22.00±2.00<sup>c</sup> | 16.00±0.00<sup>b</sup> | 0 |

Values are means of triplicate determinations; Values within the same row followed by different superscripts are significantly different at (p < 0.05).

Antimicrobial activity of antibiotic substance of isolate E is depicated on Table 1E. The inhibition zone of antibiotic substances obtained from isolate E to all standard pathogenic bacteria were significantly (P ≤ 0.05) less than the inhibition zone of VM on the same table. On the other hand, the inhibition zones of antibiotic substances (16.00±2.00) obtained from isolate E against *K. pneumonae* (multidrug resistant strain) was not sognificantly (P ≥ 0.05) different from the inhibition zone of VM (16.00±1.73). But the inhibition zones of antibiotic substances (22.00±2.00) obtained from isolate E against *MRSA* (multidrug resistant strain) was stastistically (P ≤ 0.05) greater than the inhibition zones of VM (16.00±0.00).

**Table 1E:**
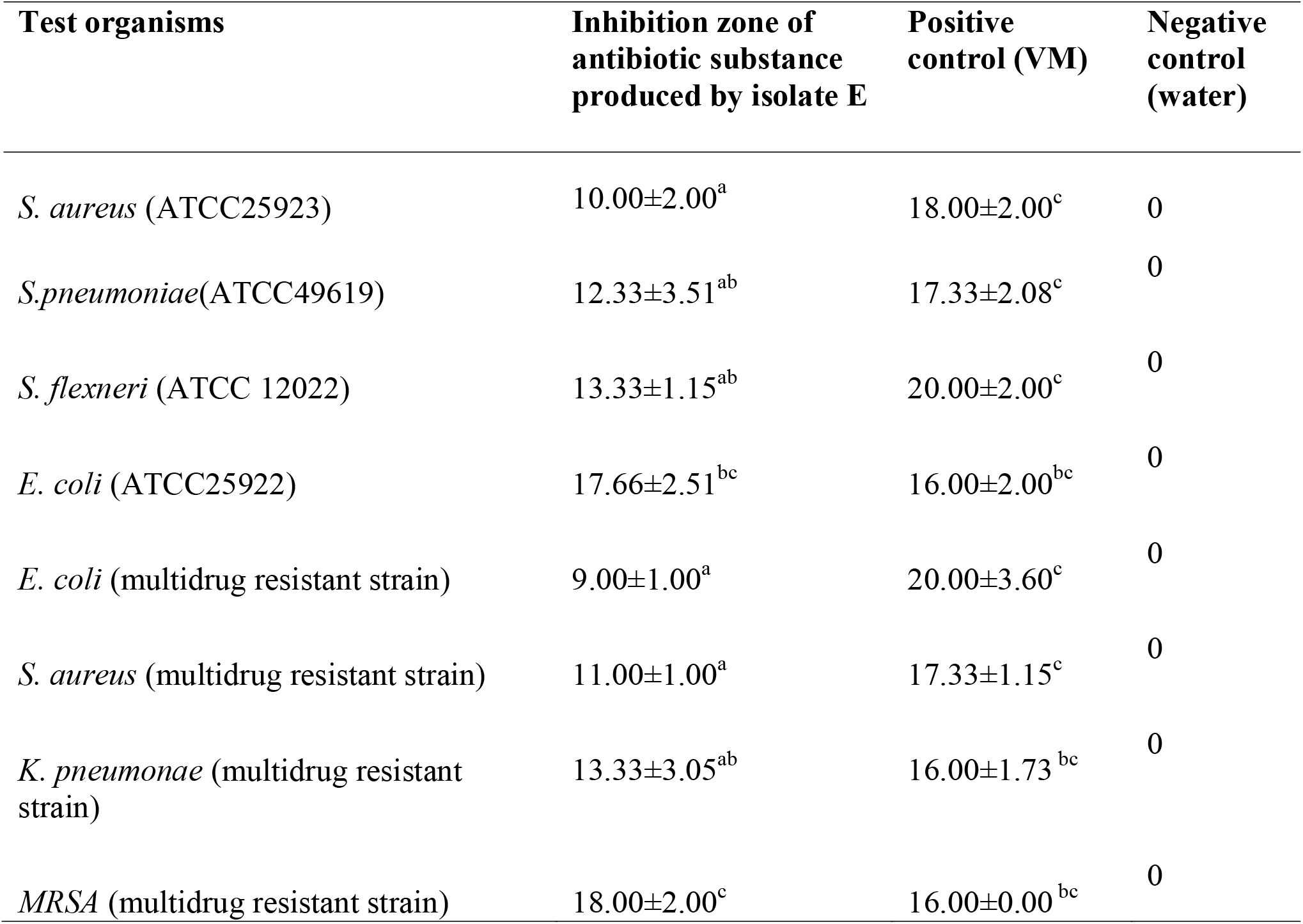
The mean inhibition zone of antibiotic substance produced by LAB against standard and drug resistant clinical pathoegnic bacteria.

Values are means of triplicate determinations; Values within the same row followed by different superscripts are significantly different at (p < 0.05).

Antimicrobial activity of antibiotic substance of isolate F is presented on Table 1F. The inhibition zone of antibiotic substances produced from isolate F against *S. aureus* (ATCC25923), *S.pneumoniae*(ATCC49619), *S. flexneri* (ATCC 12022) and *E. coli* (multidrug resistant strain) were significantly (P ≤ 0.05) less than the inhibition zone of VM. On the other hand, the inhibition zones of antibiotic substances (15.33±4.16) obtained from isolate F against *E. coli* (ATCC25922) was not stastistically (P ≥ 0.05) different from the inhibition zone of VM (16.00±2.00). Similarly, the inhibition zones of antibiotic substances (19.33±3.05) obtained from isolate F against *S. aureus* (multidrug resistant strain) was not stastistically (P ≥ 0.05) different from the inhibition zone of VM (17.33±1.15). However, the inhibition zones of antibiotic substances obtained from isolate F against *K. pneumonae* (multidrug resistant strain) and *MRSA* (multidrug resistant strain) were significantly (P ≤ 0.05) greater than the inhibition zones of VM.

**Table 1F:**
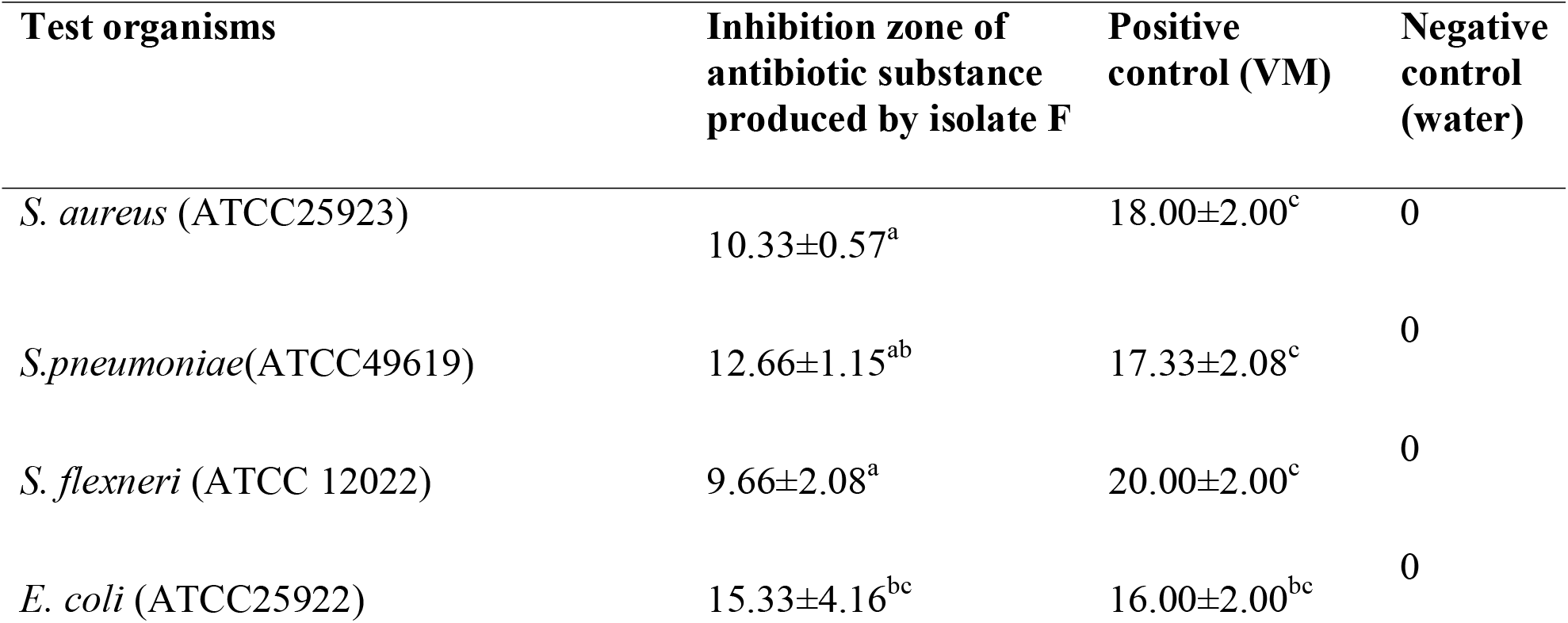
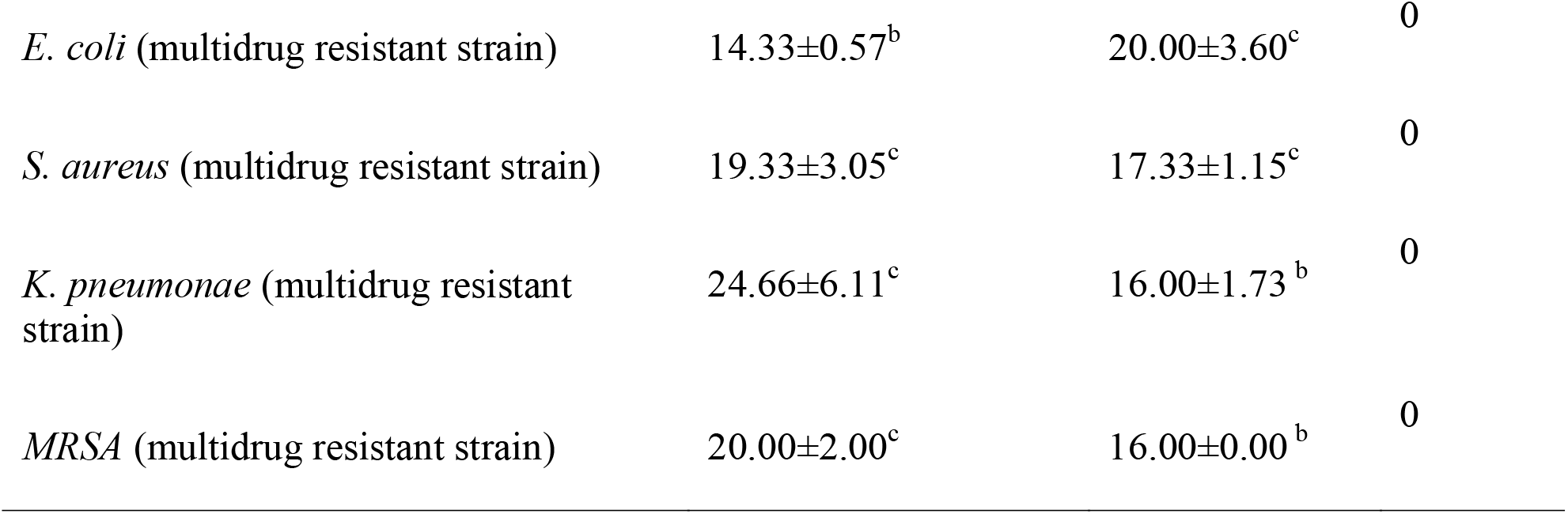
The mean inhibition zone of antibiotic substance produced by LAB against standard and drug resistant clinical pathoegnic bacteria.

| Test organisms | Inhibition zone of antibiotic substance produced by isolate F | Positive control (VM) | Negative control (water) |
| --- | --- | --- | --- |
| *S. aureus* (ATCC25923) | 10.33±0.57<sup>a</sup> | 18.00±2.00<sup>c</sup> | 0 |
| *S.pneumoniae*(ATCC49619) | 12.66±1.15<sup>ab</sup> | 17.33±2.08<sup>c</sup> | 0 |
| *S. flexneri* (ATCC 12022) | 9.66±2.08<sup>a</sup> | 20.00±2.00<sup>c</sup> | 0 |
| *E. coli* (ATCC25922) | 15.33±4.16<sup>bc</sup> | 16.00±2.00<sup>bc</sup> | 0 |
| *E. coli* (multidrug resistant strain) | 14.33±0.57<sup>b</sup> | 20.00±3.60<sup>c</sup> | 0 |
| *S. aureus* (multidrug resistant strain) | 19.33±3.05<sup>c</sup> | 17.33±1.15<sup>c</sup> | 0 |
| *K. pneumoniae* (multidrug resistant strain) | 24.66±6.11<sup>c</sup> | 16.00±1.73<sup>b</sup> | 0 |
| *MRSA* (multidrug resistant strain) | 20.00±2.00<sup>c</sup> | 16.00±0.00<sup>b</sup> | 0 |

Values are means of triplicate determinations; Values within the same row followed by different superscripts are significantly different at (p < 0.05).

Minimum inhibitory concentration (MIC) of antibiotic substances produced from isolates of LAB (A,B and C) against tested pathogenic standard and clinical multidrug resistant isolate bacteria was shown on Table 2A. Except antibiotic substance obtained from isolate C (12.5%), the MIC of isolate A and B against *S. aureus* (ATCC25923) was 6.25%. The MIC of antibiotic substance produced from isolate A,B and C against *S. pneumoniae*(ATCC49619) were 6.25%, while the MIC the MIC of isolate A against *S. flexneri* (ATCC 12022) was 6.25%. Except isolate A (12.5%), MIC of antibiotic substances produced from isolate B and C against *E. coli* (ATCC25922), *E. coli* (multidrug resistant strain) and *S. aureus* (multidrug resistant strain) was 6.25%. The MIC of antibiotic substances produced from isolate A, B and C against *K. pneumonae* (multidrug resistant strain) was 12.5%. With regard to *MRSA* (multidrug resistant strain), the MIC of antibiotic substances produced from isolate A and C against *MRSA* (multidrug resistant strain) was 6.25%, while isolate B was 12.5%.

**Table 2A:** Minimum inhibitory concentrations (MIC) of antibiotic substances produced by LAB by the dilution solutions at the dose levels of 6.25, 12.5, 25% and 50%.

| Test organisms | Solution of antibiotic substances produced by different LAB species (%) |  |  |
| --- | --- | --- | --- |
|  | A | B | C |
| *S. aureus* (ATCC25923) | 6.25% | 6.25% | 12.5% |
| *S.pneumoniae*(ATCC49619) | 6.25% | 6.25% | 6.25% |
| *S. flexneri* (ATCC 12022) | 6.25% | 12.5% | 12.5% |
| *E. coli* (ATCC25922) | 12.5% | 6.25% | 6.25% |
| *E. coli* (multidrug resistant strain) | 12.5% | 6.25% | 6.25% |
| *S. aureus* (multidrug resistant strain) | 12.5% | 6.25% | 6.25% |
| *K. pneumoniae* (multidrug resistant strain) | 12.5% | 12.5% | 12.5% |
| *MRSA* (multidrug resistant strain) | 6.25% | 12.5% | 6.25% |

Minimum inhibitory concentration (MIC) of antibiotic substances produced from isolates of LAB (D,E and F) against tested pathogenic standard and clinical multidrug resistant isolate bacteria was presented on Table 2B. Minimum inhibitory concentration (MIC) of antibiotic substances produced from isolates of LAB (D,E and F) against tested pathogenic standard and clinical multidrug resistant isolate bacteria was shown on Table 2B. Except antibiotic substance obtained from isolate E(12.5%),the MIC of isolate D and F against *S. aureus* (ATCC25923) was 6.25%.The MIC of antibiotic substance produced from isolate D and E against *S. pneumonia* (ATCC49619) were 12.5%,while the MIC of isolate F against *S. pneumonia* (ATCC49619) was 6.25%.Except isolate E(12.5%),MIC of antibiotic substance produced from isolate D and F against *S. flexneri* (ATCC 12022) were 6.25%.with regard to *E. coli* (ATCC25922),the MIC of of antibiotic substance produced from isolate E and F were 6.25%,while the MIC of antibiotic substance produced from isolate D was 25%. Except antibiotic substance produced from isolate E(6.25%),the MIC of isolate D and F against *E. coli* (multidrug resistant strain) were 12.5%. The MIC of antibiotic substance produced from isolate D,E and F against *S. aureus* (multidrug resistant strain) were 6.25%.with regard to *K. pneumonae* (multidrug resistant strain),the antibiotic substance produced from isolate D and F against *K. pneumonae* (multidrug resistant strain) were 12.5%,while isolate E was 6.25%.the MIC of antibiotic substance produced from isolate D and E against *MRSA* (multidrug resistant strain) were 6.25%,while isolate F was 12.5%. Minimum Bactericidal concentration (MBC) of antibiotic substances produced from isolates of LAB (A,B and C) against tested pathogenic standard and clinical multidrug resistant isolate bacteria was shown on Table 3A. Except antibiotic substance produced from isolate A(25%),the MBC of isolate B and C against *S. aureus* (ATCC25923) were 12.5%. with regard to *S.pneumoniae*(ATCC49619) the MBC of antibiotic substance produced from isolate A,B and C were 12.5%. the MBC of antibiotic substance produced from isolate B and C against *S. flexneri* (ATCC 12022) were 25%,while isolate A was 12.5%. %.the minimum bactericidal concentration of antibiotic substance produced from isolate A and C against *E. coli* (ATCC25922) were 25%,while antibiotic substance produced from isolate B has MBC value of 6.25%.with regard to *E. coli* (multidrug resistant strain), the MBC value of antibiotic substance produced from isolate B and C against *E. coli* (multidrug resistant strain) were 12.5%,while isolate A was 25%.Except antibiotic substance produced from isolate C(6.25%),MBC value of antibiotic substance produced from isolate A and B were 12.5% against *S. aureus* (multidrug resistant strain. antibiotic substance produced from isolate A,B and C have MBC value of 25%,50% and 12.5% respectively against *K. pneumonae* (multidrug resistant strain).with regard to *MRSA* (multidrug resistant strain),MBC of antibiotic substance produced from isolate A and C were 25%,while isolate B was 12.5%.

**Table 2B:** Minimum inhibitory concentrations (MIC) of antibiotic substances produced by LAB by the dilution solutions at the dose levels of 6.25, 12.5,25% and 50%.

|  | **D** | **E** | **F** |
| --- | --- | --- | --- |
| *S. aureus* (ATCC25923) | 6.25% | 12.5% | 6.25% |
| *S.pneumoniae*(ATCC49619) | 12.5% | 12.5% | 6.25% |
| *S. flexneri* (ATCC 12022) | 6.25% | 12.5% | 6.25% |
| *E. coli* (ATCC25922) | 25% | 6.25% | 6.25% |
| *E. coli* (multidrug resistant strain) | 12.5% | 6.25% | 12.5% |
| *S. aureus* (multidrug resistant strain) | 6.25% | 6.25% | 6.25% |
| *K. pneumoniae* (multidrug resistant strain) | 12.5% | 6.25% | 12.5% |
| *MRSA* (multidrug resistant strain) | 6.25% | 6.25% | 12.5% |

**Table 3A:** Minimum bactericidal concentrations (MBC) of antibiotic substances produced by LAB by the dilution solutions at the dose levels of 6.25, 12.5, 25% and 50%.

| Test organisms | Solution of antibiotic substances produced by different LAB species (%) |  |  |
| --- | --- | --- | --- |
|  | A | B | C |
| *S. aureus* (ATCC25923) | 25% | 12.5% | 12.5% |
| *S.pneumoniae*(ATCC49619) | 12.5% | 12.5% | 12.5% |
| *S. flexneri* (ATCC 12022) | 12.5% | 25% | 25% |
| *E. coli* (ATCC25922) | 25% | 6.25% | 25% |
| *E. coli* (multidrug resistant strain) | 25% | 12.5% | 12.5% |
| *S. aureus* (multidrug resistant strain) | 12.5% | 12.5% | 6.25% |
| *K. pneumoniae* (multidrug resistant strain) | 25% | 25% | 12.5% |
| *MRSA* (multidrug resistant strain) | 25% | 12.5% | 25% |

Minimum inhibitory concentration (MIC) of antibiotic substances produced from isolates of LAB (D,E and F) against tested pathogenic standard and clinical multidrug resistant isolate bacteria was shown on Table 3B.Except antibiotic substance obtained from isolate E(12.5%),the MIC of isolate D and F against *S. aureus* (ATCC25923) was 6.25%.The MIC of antibiotic substance produced from isolate D and E against *S. pneumoniae*(ATCC49619) were 12.5%,while the MIC of isolate F against *S. pneumoniae*(ATCC49619) was 6.25%.Except isolate E(12.5%),MIC of antibiotic substance produced from isolate D and F against *S. flexneri (*ATCC 12022) were 6.25%.with regard to *E. coli* (ATCC25922),the MIC of of antibiotic substance produced from isolate E and F were 6.25%,while the MIC of antibiotic substance produced from isolate D was 25%. Except antibiotic substance produced from isolate E(6.25%),the MIC of isolate D and F against *E. coli* (multidrug resistant strain) were 12.5%. The MIC of antibiotic substance produced from isolate D,E and F against *S. aureus* (multidrug resistant strain) were 6.25%.with regard to *K. pneumonae*(multidrug resistant strain),the antibiotic substance produced from isolate D and F against *K.pneumonae* (multidrug resistant strain) were 12.5%,while isolate E was 6.25%.the MIC of antibiotic substance produced from isolate D and E against *MRSA* (multidrug resistant strain) were 6.25%,while isolate F was 12.5%.

**Table 3B:** Minimum bactericidal concentrations (MBC) of antibiotic substances produced by LAB by the dilution solutions at the dose levels of 6.25, 12.5,25% and 50%.

| Test organisms | Solution of antibiotic substances produced by different LAB species (%) |  |  |
| --- | --- | --- | --- |
|  | D | E | F |
| *S. aureus* (ATCC25923) | 25% | 25% | 12.5% |
| *S.pneumoniae*(ATCC49619) | 12.5% | 12.5% | 25% |
| *S. flexneri* (ATCC 12022) | 6.25% | 25% | 25% |
| *E. coli* (ATCC25922) | 25% | 12.5% | 25% |
| *E. coli* (multidrug resistant strain) | 12.5% | 12.5% | 25% |
| *S. aureus* (multidrug resistant strain) | 25% | 25% | 12.5% |
| *K. pneumoniae* (multidrug resistant strain) | 12.5% | 12.5% | 25% |
| *MRSA* (multidrug resistant strain) | 12.5% | 25% | 25% |

Minimum Bactericidal concentration (MBC) of antibiotic substances produced from isolates of LAB (A,B and C) against tested pathogenic standard and clinical multidrug resistant isolate bacteria was shown on Table 3A.Except antibiotic substance produced from isolate A(25%),the MBC of isolate B and C against *S. aureus* (ATCC25923) were 12.5%. with regard to *S.pneumoniae*(ATCC49619) the MBC of antibiotic substance produced from isolate A,B and C were 12.5%. the MBC of antibiotic substance produced from isolate B and C against *S. flexneri* (ATCC 12022) were 25%,while isolate A was 12.5%. %.The minimum bactericidal concentration of antibiotic substance produced from isolate A and C against *E. coli* (ATCC25922) were 25%,while antibiotic substance produced from isolate B has MBC value of 6.25%. With regard to *E. coli* (multidrug resistant strain), the MBC value of antibiotic substance produced from isolate B and C against *E. coli* (multidrug resistant strain) were 12.5%,while isolate A was 25%. Except antibiotic substance produced from isolate C(6.25%),MBC value of antibiotic substance produced from isolate A and B were 12.5% against *S. aureus* (multidrug resistant strain. Antibiotic substance produced from isolate A,B and C have MBC value of 25%, 50% and 12.5% respectively against *K. pneumonae* (multidrug resistant strain). With regard to *MRSA* (multidrug resistant strain),MBC of antibiotic substance produced from isolate A and C were 25%,while isolate B was 12.5%.

Minimum Bactericidal concentration (MBC) of antibiotic substances produced from isolates of LAB (D,E and F) against tested pathogenic standard and clinical multidrug resistant isolate bacteria was shown on Table 3B. MBC of antibiotic substance produced from isolate D against S.pneumoniae(ATCC49619), *E. coli* (multidrug resistant strain), *K. pneumonae*(multidrug resistant strain) was 12.5%,while MBC of isolate D against *E. coli* (ATCC25922) and S*. aureus* (multidrug resistant strain) was 25%. MBC of antibiotic substance produced from isolate D against *S. aureus* (ATCC25923) and *S. flexneri* (ATCC 12022) were 25% and 6.25% respectively. With regard to antibiotic substance produced from isolate E, the MBC against *S.pneumoniae*(ATCC49619), *E. coli* (ATCC25922), E. coli (multidrug resistant strain) and *K. pneumonae* (multidrug resistant strain) were 12.5%, while MBC of isolate E against *S. flexneri* (ATCC 12022), *S. aureus* (multidrug resistant strain) and *MRSA* (multidrug resistant strain) was 25%.isolate E’s highest MBC value recorded was against *S. aureus* (ATCC25923)(25%). MBC of antibiotic substance produced from isolate F against *S.pneumoniae* (ATCC49619), *E. coli* (ATCC25922), *E. coli* (multidrug resistant strain) and *MRSA* was 25%, while MBC of isolate F against *S. aureus* (ATCC25923) and *S. aureus* (multidrug resistant strain) was 12.5%.

Minimum inhibitory concentration of different concentration of antibiotic substances produced from diffent isolates of metata LAB are shown in Table 3A and 3B. MIC of antibiotic substances produced from diffent isolates of metata LAB at 6.25% against total test pathogenic bacteria was 62.5%. On the other hand, MIC of antibiotic substances produced from diffent isolates of metata LAB at 6.25% against multidrug resistant clinical test pathogenic bacteria was 54.2%. MIC of antibiotic substances produced from diffent isolates of metata LAB at 6.25% against Gram positive and negative were 70.8% and 45.8%, respectively.

## 5. Discussion

Emerging and re-emerging of human pathogens are becoming the current threat worldwide. Thus, finding new antibacterial agents for the treatment of pathogens from microorganisms like LAB is a basic and important scenario (Assefa, e al, 2008). LAB display numerous antimicrobial activities in fermented foods. Moreover, the antimicrobial activity of metabolites produced from antibiotic producing microorganisms can be influenced by the type of the pathogen used, the physical and chemical parameters used under different environmental conditions (Bizuye, et al., 2013). In this study, antibiotic substance produced from LAB in metata ayib showed antibacterial activities against standard and drug resistant tested micro organisms.The antibiotic substance test against the test organisms in the agar well diffusion method and the MIC,MBC confirm the potential antibacterial activity of the isolates in the study. Now a day’s infections caused by bacteria become serious problem(Odimegwu et al., 2008). Some of the common bacterial pathogen included in this study were S.aureus (a wound infecting pathogen which can cause septicemia), *S.pneumoniae* (cause for lung inflammation), E.coli (a most common bacteria of which virulent strains can cause gastroenteritis, urinary tract infections, neonatal meningitis), *K.pneumoniae* (which is the causative organism of pneumonia)(Juthani-mehta, 2007).

The inhibition zone of the antibiotic substance against the standard and drug resistant test organisms in this study ranged from 9mm up to 25.33mm however, In the previous studies the inhibition zone of the supernatant from LAB isolates against standard human pathogens was ranged from 8 mm to 10 mm (Savadogo, 2004). and the inhibition zone of supernatant of LAB against human pathogens was ranged from 9.25 ± 0.35mm to 11.8 ± 0.35 mm(Assefa et al., 2008). The result of this study shows greater inhibition zone than earlier studies. This might be due to the production of organic acids, but also other compounds, such as ethanol, hydrogen peroxide, diacetyl, reuterin and bacteriocins. The therapeutic effect of the antibiotic substance from Metata ayib has a promising antimicrobial activity and better to processed in pharmaceutical companies to be prepared as an alternative medicinal drug against the infections caused by common pathogenic bacteria studied in this research.

According to Saranya and Hemashenpagam (2011) report, the inhibition zone of cell free supernatant against clinical strains of *E. coli, K. pneumonia and S. aureus* was ranged from 6 to14 mm, 7 to 14 mm, and 6 to13 mm, respectively.(Hemashenpagam and Saranya, 2011). The present study showed that the inhibition zone against clinical strains of *E.coli, K.pneumonia, and S.aureus*(ATCC25923) was ranged 8.65±0.58, 10.11± 3.01, and 14.67±2.52, respectively. As indicated here, the present study result showed better inhibition zone against *E. coli, K. pneumonia and S. aureus* when compared to Saranya and Hemashenpagam(2011). The result of this study showed that all the standard and drug resistance isolates of bacterial strains were susceptible. In addition most the tested bacterial strains showed MIC value 6.25% against the antibiotic substance. Therefore the antibiotic substance will serve as an effective way of controlling microbial infections especially caused by those drug resistant pathogens. Inhibition variety of pathogenic bacteria by LAB due to a combination of many factors like production of lactic acid which reduce pH of *Metata Ayib* and also other inhibitory substances such as bacteriocins which are responsible for the most antimicrobial activity.

According to this study, The MICs and MBCs were determined against the tested bacterial strains that showed susceptibility to the antibiotic isolate. The MIC of antibiotic substance produced from *Metata Ayib* against for both tested standard and clinical isolate bacteria were ranged from 6.25-12.5 %. The MBC of the antibiotic substance produced frmo *Metata ayib* against for both tested standard and clinical isolates were ranged from 6.25%-50%.Similar results of the MIC and MBC value were reported by (Beyan et al., 2011). From this study, it is possible to suggest that antibiotic substances produced from Metata Ayib can inhibit bacteria growth and thereby it can play a role in the antibacterial medicinal investigations. In the present study, there were few results which are not parallel among methods used to determine antimicrobial activity (agar well diffusion and serial dilution).In the agar well diffusion assay few antibiotic isolates slightly showed better antibacterial activities against drug resistant isolates than the standard strains. Its novel to take consideration those clinical isolates were resistant to the commercial antibiotic vancomycin,on the other outcomes, growth of standard strains were inhibited at low concentration (6.25%) than drug resistant isolates during serial dilution method. This can strengthen some reports (Youssef and Tawil,1980) showed that comparison of methodologies (agar well diffusion and serial dilution)which are used to evaluate the antimicrobial activity may not usually give compatible results. Antibiotic substances was considered as having antimicrobial activity if it inhibits a particular organism at 50% concentration in the agar well diffusion assay(Parekh et al., 2005). In this study, it was promising that antibiotic substances have an effective antimicrobial activity.Therefor its hopeful that antibiotic substance from LAB in Metata ayib have antimicrobial potential to inhibit growth of tested pathogens, especially if further processed in to new drugs.

The antibiotic substance was significantly inhibited gram positive bacteria than gram negative isolates which shares the clam’s that gram negative bacteria are more resistant due to the presence of their outer wall containing lipopolysaccharide which make the cell wall impermeable to lipophilic solutes. The overall antimicrobial activity of the antibiotic substance to both gram positive and gram negative bacterial strains, especially to those drug resistant clinical isolates, reveals that the therapeutic potentials of the antibiotic substance can be used along with pure drugs to control continually emerging resistant bacteria, which are becoming a threat to human health.

Research on antimicrobial substances produced by LAB, has led to their potential use as natural preservatives. These antibacterial compounds may be used to combat the growth of pathogenic microorganisms in the food industry. Antimicrobial compounds can be applied either as purified chemical agents or as viable cultures in the case of fermented products(Barnby-Smith, 1992). Novel purified antimicrobial compounds require convincing data to substantiate their lack of toxicity to ensure their acceptance before use in various food processes.

## 6. CONCLUSION AND RECOMMENDATION

### 6.1. Conclusion

In this investigation, the culture filtrates of LAB isolated from metata ayib have been exhibited antimicrobial activity against pathogenic test bacterial strains. Lactic acid becteria isolates have a strong antimicrobial activity against pathogenic and food spoilage microorganisms and play a vital role in the preservation of metata ayib. The possible reason of LAB isolates, which capable of inhibiting the growth of pathogenic bacteria, was maybe due to production of lactic acid, acetic acid and secondary metabolites like bacteriocins.

From the overall study it canbe concuded that, LAB from Metata ayib can produce different metabolites having antimicrobial and probiotic activities. Cultivation of these potential isolates under optimum condition can lead for the production of potential antibacterial agents (like lactic acid, hydrogen peroxide and others) having preservation as well as probiotic activity. These can reduce and control different human health problems caused by pathogens. Therefore, isolation and screening of LAB from potential locally prepared Metata Ayib is the basic sources for the discovery of new potential LAB for controlling and treatment of infectious disease to improve the health quality of human beings.

In general, the finding of the present study shows that the presence of the LAB in the Metata Ayib help to avoid the growth of other pathogenic microorganism and production of antibiotics which can exhibit antimicrobial activity against some common microorganism which can cause diseases. LAB display numerous antimicrobial activities in fermented foods. This is mainly due to the production of organic acids, but also of other compounds, such as ethanol, hydrogn peroxide, diacetyl, reuterin and bacteriocins. This study underline the important role of bacteriocinogenic LAB that may play in food industry as starter cultures to improve food quality and safety.

The results of this investigation can also provide baseline for information for future studies about the antibacterial activity of LAB from fermented foods like Metata ayib. Generally the outcome of this research implies that antibiotic substance from Metata ayib contain compounds which have antibacterial activities and its important to conduct additional research studies on Metata Ayib.

### 6.2. Recommendations

Based on the finding of the present study, the following recommendation can be forwarded:

➢ An *in vitro* investigation of antibiotic substance from Metata Ayib studied in this research demonstrated an inhibitory effect towards all tested pathogenic bacteria. Therefore, further *in vivio* tests is recommended to evaluate therapeutic capability of Metata Ayib in detail.
➢ The community should be aware of the scientific acceptance of therapeutic potentials of Metata Ayib with the indigenous knowledge. This is because the society as traditional level use metata ayibe for treatment of various infectious diseases.
➢ Additional scientific researches and investigations need to be conducted to understand the potential application of the spices which are incorporated as ingredient during the Metata Ayib preparation.

## 8. APPENDIX

A,Inhibition zone of Antibiotic substance isolate A against MRSA

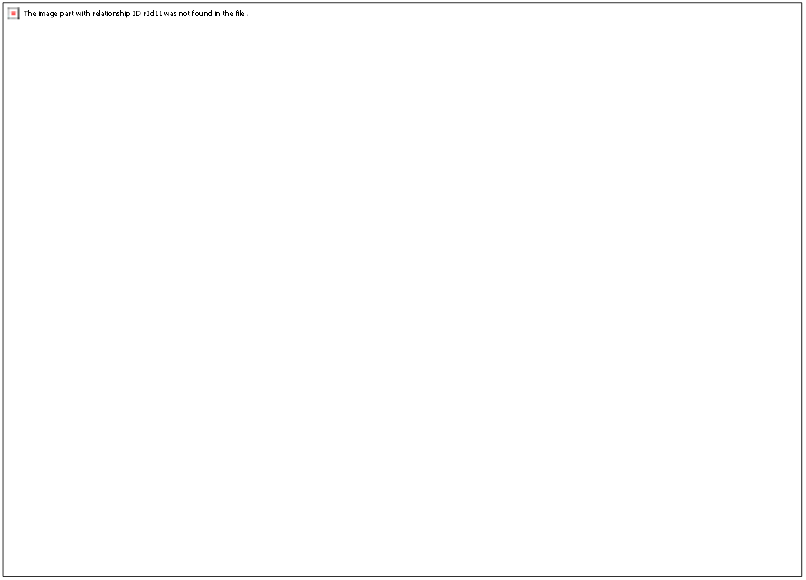

B, Inhibiotion zone of antibiotic isolate A against *E.coli*

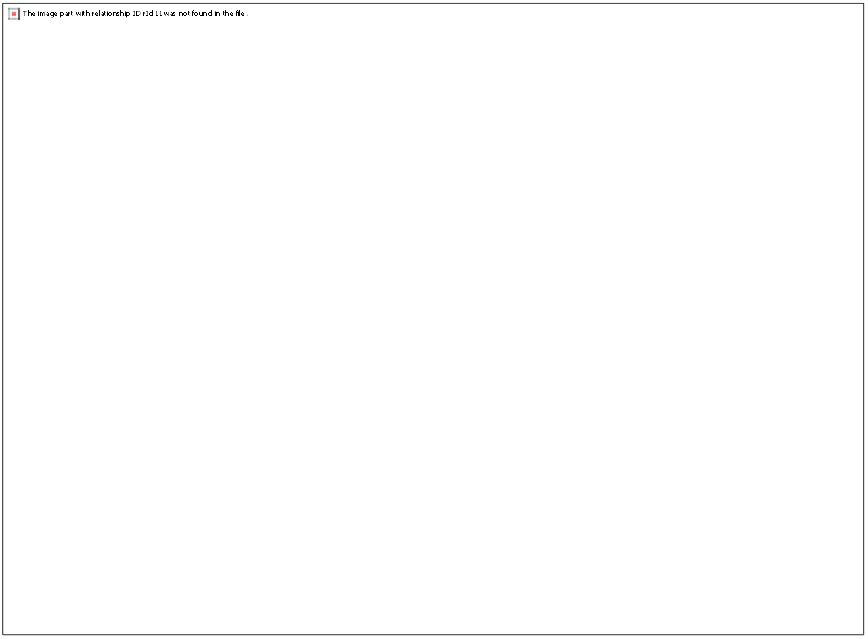

C,Inhibition zone of antibiotic substance isolate E against *S. aureus* (ATCC25923)

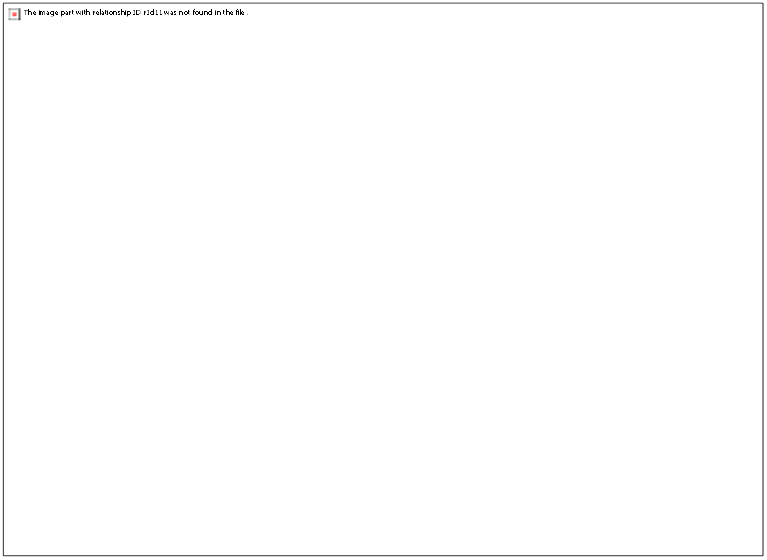

## REFERENCES

Anteneh, T., Tetemke, M., Mogessie, A., 2011. Antagonism of lactic acid bacteria against foodborne pathogens during fermentation and storage of borde and shamita, traditional Ethiopian fermented beverages. Int. Food Res. J. 18.

Ashenafi, M., Beyene, F., 1993. Effect of container smoking and udder cleaning on the microflora and keeping quality of raw milk from a dairy farm in Awassa.

Ashenafi, M., Mehari, T., 1995. Some microbiological and nutritional properties Some microbiological and nutritional properties of Borde and Shamita, traditional Ethiopian fermented beverages. Ethiop. J. Heal. Dev. 9, 105–110.

Assefa, E., Beyene, F., Santhanam, A., 2008. Effect of temperature and pH on the antimicrobial activity of inhibitory substances produced by lactic acid bacteria isolated from Ergo, an Ethiopian traditional fermented milk. African J. Microbiol. Res. 2, 229–234.

Barnby-Smith, F.M., 1992. Bacteriocins: applications in food preservation. Trends Food Sci. Technol. 3, 133–137. https://doi.org/10.1016/0924-2244(92)90166-T

Beresford, T.P., Fitzsimons, N.A., Brennan, N.L., Cogan, T.M., 2001. Recent advances in cheese microbiology. Int. Dairy J. 11, 259–274. https://doi.org/10.1016/S0958-6946(01)00056-5

Beyan, A., Ketema, T., Bacha, K., 2011. Antimicrobial susceptibility pattern of lactic acid bacteria isolated from ergo, a traditional Ethiopian fermented milk, Jimma, south west Ethiopia. Ethiop. J. Educ. Sci.

Bizuye, A., Moges, F., Andualem, B., 2013. Isolation and screening of antibiotic producing actinomycetes from soils in Gondar town, North West Ethiopia. Asian Pacific J. Trop. Dis. 3, 375–381. https://doi.org/10.1016/S2222-1808(13)60087-0

Chelule, P.K., Mokoena, M.P., Gqaleni, N., 2010. Advantages of traditional lactic acid bacteria fermentation of food in Africa 1160–1167.

Daeschel, M.A., 1993. Applications and Interactions of Bacteriocins from Lactic Acid Bacteria in Foods and Beverages, Bacteriocins of Lactic Acid Bacteria. ACADEMIC PRESS, INC. https://doi.org/10.1016/b978-0-12-355510-6.50012-9

Das, A.J., Deka, S.C., 2012. Fermented foods and beverages of the North-East India. Int. Food Res. J. 19, 377–392.

Egerva, M., Lindmark, H., Roos, S., Huys, G., Lindgren, S., 2007. Effects of Inoculum Size and Incubation Time on Broth Microdilution Susceptibility Testing of Lactic Acid Bacteria 51, 394–396. https://doi.org/10.1128/AAC.00637-06

Gebreselassie, N., Abrahamsen, R.K., Beyene, F., Abay, F., Narvhus, J.A., 2016. Chemical composition of naturally fermented buttermilk. Int. J. Dairy Technol. 69, 200–208. https://doi.org/10.1111/1471-0307.12236

Gonfa, A., Foster, H.A., Holzapfel, W.H., 2001. Field survey and literature review on traditional fermented milk products of Ethiopia. Int. J. Food Microbiol. 68, 173–186. https://doi.org/10.1016/S0168-1605(01)00492-5

Hemashenpagam, N., Saranya, S., 2011. Antagonistic activity and antibiotic sensitivity of Lactic acid bacteria from fermented dairy products. Adv. Appl. Sci. Res. 2, 528–534.

Holzapfel, W.H., Haberer, P., Geisen, R., Schillinger, U., 2001. Taxonomy and important features of probiotic microorganisms in PHYSIOLOGIC PROPERTIES OF LACTIC ACID BACTERIA. Am. J. Clin. Nutr. https://doi.org/10.1093/ajcn/73.2.365s

Israti, D., et al, 2011. Received 29 September 2011 Revised 10 October 2011 Fresh beef slices were marinated by immersion in marinades based on dry red wine, lime-tree honey, salt, spices and seasoning plants as thyme (35, 75–85.

Jay, J.M., 1998. Fermentation and Fermented Dairy Products 131–148. https://doi.org/10.1007/978-1-4615-7476-7_7

Juthani-mehta, M., 2007. Asymptomatic Bacteriuria and Urinary Tract Infection in Older Adults 23, 585–594. https://doi.org/10.1016/j.cger.2007.03.001

O’mahony, M., Thieme, U., Goldstein, L.R., 1988. The Warm-up Effect as a Means of Increasing the Discriminability of Sensory Difference Tests. J. Food Sci. 53, 1848–1850. https://doi.org/10.1111/j.1365-2621.1988.tb07858.x

Odimegwu, D., Ibezim, E.C., Okoye, F., 2008. Wound healing and antibacterial activities of the extract of Dissotis theifolia (Melastomataceae) stem formulated in a simple ointment base. J. Med. Plants Res. 2, 011–016.

Parekh, J., Chanda, S., n.d. Antibacterial and phytochemical studies on twelve species of Indian medicinal plants 10, 175–181.

Seifu, E., 2013. Chemical composition and microbiological quality of Metata Ayib: A traditional Ethiopian fermented cottage cheese. Int. Food Res. J. 20, 93–97.

Stiles, M.E., Holzapfel, W.H., 1997. Lactic acid bacteria of foods and their current taxonomy. Int. J. Food Microbiol. 36, 1–29. https://doi.org/10.1016/S0168-1605(96)01233-0

Tannock, G.W., n.d. Probioticproperties of lactic-acid bacteria: plentyof scopeI fundamental R & D for.

Yvon, M., Rijnen, L., 2001. Cheese flavour formation by amino acid catabolism. Int. Dairy J. 11, 185–201. https://doi.org/10.1016/S0958-6946(01)00049-8

